# Biomathematical model of prebiotic autocatalytic RNA networks shows birth of adaptive immunity from degenerating molecular parasite catalysts

**DOI:** 10.1101/2023.05.25.542273

**Authors:** Magnus Pirovino, Christian Iseli, Joseph A. Curran, Bernard Conrad

## Abstract

Catalysis and specifically autocatalysis are the quintessential building blocks of life. However, autocatalytic networks are necessary but not sufficient for life-like properties such as self-propagation and adaptation to emerge. We previously showed using Lotka-Volterra equations that a tripartite, early life cycle comprising a host-template, molecular parasites and hyperparasites (parasites of parasites) conferred superior stability to the habitat. Here we build on this model, and on the seminal notion that chemical networks and ecologically interacting biological species are akin, implying an instructive equational analogy between catalysis and ecology, both accurately modelled by equations from the Lotka-Volterra family. Template-based autocatalytic RNA networks seed the spontaneous emergence of immunity against molecular parasites, with degenerating molecular parasite catalysts being the founding seed of the homeostatic process. First, upon molecular parasite encounter, specific ribocatalysts providing parasite resistance appear in primordial autocatalytic cycles, supplanted by promiscuous, efficiently self-replicating, but parasite sensitive ribozymes. This is supported by catalyst tradeoff analysis: substrate promiscuity confers high catalytic activity, but entails parasite exposure, higher substrate specificity implies relatively lower activity, yet it offers parasite resistance. Second, sustained parasite influx generates hyperparasite cycles, memorizing parasite-specific catalyst-subunits that now once more provide parasite resistance, in a homeostatically stabilized habitat. Third, under continuous exposure to progressively elaborate parasite populations, hyperparasite catalysts degenerate and embody antiparasite immunity. The observed triggering of microbial immunity by hyperparasitic microbial genomes are in agreement with this model. As such, it offers an attractive and unifying theory for the spontaneous birth of immunity in early life.

**Author Summary:** The quintessential components of life comprise a potent acceleration of naturally occurring, but improbable chemical reactions (catalysis), and the arrangement of accelerated chemical reactions in closed loops (autocatalytic sets). This is required, but is not sufficient for such networks to self-propagate and adapt. We previously developed an early life model for a stabilized habitat of networks, using a family of equations shown to accurately model both chemical reactions and the interactions of biological species. This model became stabilized only if the molecular host species was parasitized, an event considered unavoidable, and at the same time, the parasite itself was also parasitized. We now built on that model, using the same family of equations, by letting individual autocatalytic sets interact. The new results show that primitive autocatalytic networks emerge spontaneously, but inefficiently and resist parasites. With time, they become more efficient, which renders them parasite-sensitive. Upon continuous external parasite invasion, the primitive, parasite-resistant catalysts (hyperparasites) that still persist are recruited, conferring stability to the habitat. If the habitat is further overwhelmed with parasites, it selects the elegant solution of letting the parasite-resistant catalysts degenerate, which now produces immunity to those parasites (“antibodies”). This represents the birth of adaptive immunity.

## Introduction

The multistep transition process from inanimate to living systems has been dated to >3.9 Ga [1]. A non-exhaustive list of thresholds that needed to be progressively surmounted during this process has been drawn, comprising, among others, chirality symmetry breaking, spontaneous polymerization, self-assembly of compartments, and particularly the catalytic closure threshold [2]. While autocatalysis is the core ingredient of life, it is not sufficient. The ability of prebiotic networks to self-propagate and adapt hinges upon multiple, sequential interactions of autocatalytic subsystems among themselves and with the environment, creating higher order structures [3-7]. In one of those models, pairs of autocatalytic cycles following chemical mass action kinetics [8] exhibit the superior complexity-building behaviors usually attributed to biological species, for instance competitive, predator-prey or mutualistic relationships [9]. Predator-prey type autocatalytic cycles display a dynamics similar to the Lotka-Volterra (LV) model [9].

One central pillar in the theory of how early life could have materialized pertains to the “RNA world”. This stipulates that the initial informational polymer was RNA, and that the same molecule was also the fundamental catalyst [10]. The underlying concept of autocatalytic sets was laid out more than half a century ago [11,12]. It was subsequently refined [3] and validated in a proof of principle experiment showing that RNA fragments could build self-replicating ribozymes from individual RNA fragments via cooperative catalytic cycles [13]. In parallel, an RNA polymerase ribozyme was generated by *in vitro* evolution that exhibited a high level of processivity [14]. Collectively, both theoretical considerations and experimental evidence support the view of a gradual evolution towards RNA self-catalysis. This started with self-assembly, followed by the formation of autocatalytic networks using self-assembled substrates. From this emerged different ribozymes (recombinases, ligases) and template based ligation, leading to a template based, self-replicase catalyzed assembly of monomers [15].

RNA self-catalysis is unthinkable without access to generic energy levels. It is commonly argued that autocatalytic chemical networks preceded template-based autocatalysis [16]. Indeed, proto-metabolic reactions without metals and enzymes have been demonstrated, suggesting the existence of an ancestral analogue of the core reverse Krebs cycle [17]. By force of circumstances, autocatalytic metabolic and template-based networks are bound to have joined forces to render the emergence of protocells sufficiently likely [18].

The ribozyme catalyzed RNA-polymerization process is inherently error prone [10,14], generating variant sequence space mutants called a quasispecies [19,20]. This variant sequence-space intrinsically implies the obligate presence and evolutionary persistence of the molecular parasites associated with all life forms, as argued essentially for thermodynamic reasons [21,22]. We previously developed a prebiotic hyperparasite framework based on the LV-equations [23], demonstrating that superior homeostasis arose from the tripartite configuration within a given molecular habitat, namely, the molecular host, the parasite and the hyperparasite, i.e. the parasite of the parasite. The empirical evidence of highly prevalent tripartite communities in the virosphere lends substantial credit to this notion [24-27]. The LV-equations have become the default tool to quantitatively assess interactions in microbial communities [28]. However, it is often forgotten that Alfred Lotka initially modelled catalysis and autocatalysis [29]. Based on his equations modeling chemical reactions that transform a substrate [*S*] into a product [*P*] via two intermediates, *X* and *Y*, if the reactions producing *X* and *Y* are assumed to be autocatalytic, then the resulting ordinary differential equation is the classical LV-predator-prey system [30]. This stoichiometry corresponds to the Michaelis and Menten mechanism - the universal formalism for catalysis - that considers fully reversible reactions *S* + *E* ⇌ *B1* ⇌ *B2* ⇌ *P* + *E*; (*B1* and *B2* are the intermediates equivalent to *X* and *Y*) [31]. In fact, LV-equations commonly used in ecology, the Rosenzweig-MacArthur model with Holling type II response [32,33], are analogous to the Michaelis-Menten mechanism in enzyme kinetics [34]. Jointly, these considerations lay the ground for modelling catalysis, autocatalysis and interactions within primordial, template-based ecological RNA communities with one single biomathematical tool, namely the LV-model. This reasoning has led us to extend the initial tripartite model, building on multiple, sequentially interacting autocatalytic RNA networks memorizing the historical parasite exposure. We model, how molecular parasite resistance and antiparasite immunity can be selected for and gradually emerge from progressively degenerating hyperparasites in an intrinsic fashion in early life, in hosts exposed to increasingly more complexed parasite populations.

## Results

In a stepwise and gradual process described in the Supplementary Information (proofs, Suppl. Inf.) and Methods sections, we lay out using the LV-equations, how simple template-based RNA autocatalytic cycles are initially built and how over time they become increasingly more elaborated as a result of molecular parasite exposure. Metabolic autocatalytic networks are assumed to be present and to provide sufficient energy [23]. The coexistence of prebiotic ecology/heredity [35,36] with an emerging canonical, template-based Darwinian heredity allows for the complexification of the quasispecies that in turn reflects its historical parasite exposure. The previously established tripartite habitat framework featuring a template-based RNA host, a molecular parasite that is itself parasitized (hyperparasite; [23]), is instrumental for homeostasis and the emergence of resistance, and finally immunity to the increasingly more exhaustive molecular parasite populations.

### Autocatalytic cycles in transitional early RNA-settings

The seminal duality of the RNA world, namely the fact that the same RNA molecule provides both the template and the self-replicating catalyst [10] constitutes the foundation of the initial autocatalytic cycles (replication equations (1.1), (1.2), forming a two-step autocatalytic cycle; proof outlined in Suppl. Inf.). Since the template and the enzyme/ribozyme remain in principle unchanged during this process, both can be viewed as catalysts, as long as the molecules do not undergo deleterious loss of function mutations.

### Molecular parasitism in autocatalytic cycles

Molecular parasites are an inherent feature of the replication process, they are inseparably associated with all life forms [21,22], for a number of reasons, the most important being the accuracy-rate tradeoff [37,38]. Average enzymes operate orders of magnitude (k_cat_/K_M_ of ∼10^5^ s^−1^ M^−1^) below the diffusion rate (10^8^ -10^9^ s^−1^ M^−1^), which implies promiscuous substrate binding, high catalytic activity, and as a secondary tradeoff a high mutation rate with a large variance in the quasispecies (speed being more important than enzyme accuracy; [39]). In an initial autocatalytic cycle, self-replicating ribozymes emerge (ribozyme *R*_2_ acting on a replicating template *R*_1_; proof outline in Suppl. Inf.; Model 1^(1)^ in Methods). The catalyst tradeoff opposes specific, but catalytically relatively less active molecules, and promiscuous, yet catalytically highly efficient molecules [37,38]. Therefore, the catalytically very efficient and promiscuous *R*_2_ will inevitably be parasitized by RNA templates *P*. Figure 1 illustrates the population sizes of the habitat constituents under high (Fig. 1A), and low (Fig. 1B) parasite exposure. As expected, high parasite loads overwhelm the habitat with *P* templates an observation that begs a solution.

**Figure 1.**
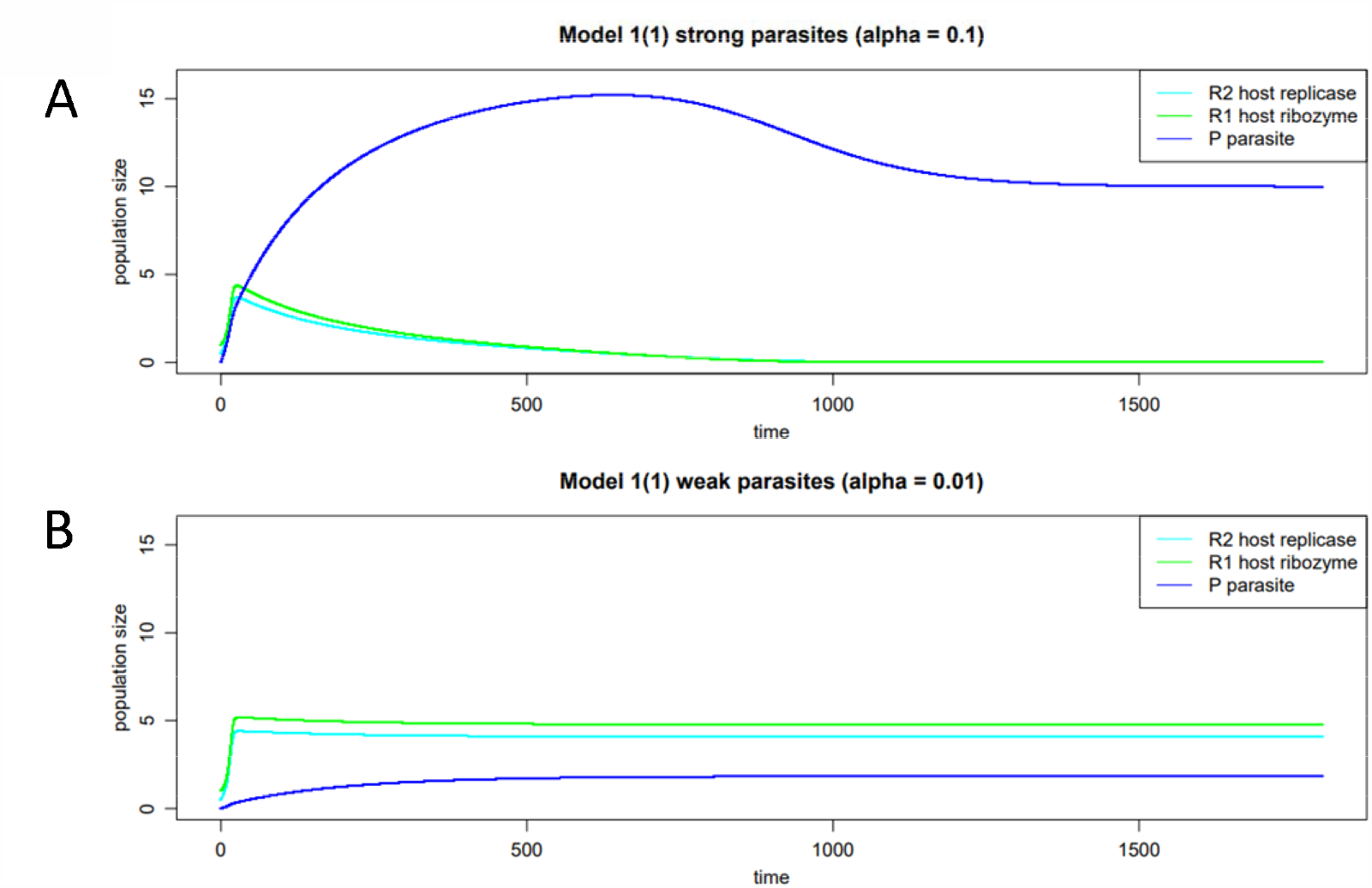
Habitat with one autocatalytic host cycle, variably parasite exposed. Fig. 1A. The catalysts in an autocatalytic cycle (ribozyme *R*_2_, template *R*_1_) exposed to incoming parasites *P* are promiscuous and hence also amplify *P*. Under high parasite exposure, parasites will therefore outcompete host molecules. Fig. 1B. Under weak parasite exposure, the host molecules *R*_1_ and *R*_2_ prevail.

### A stable, primordial autocatalytic cycle in the presence of molecular parasites

In a primordial habitat with abundant inflow of molecular RNA species including parasites, a quasispecies will eventually emerge with the central feature that it also comprises molecules with superior parasite-specificity, which implies relatively lower catalytic activity, and confers parasite resistance (protoreplicase *E*_2,_ prototemplate *F*_1_; equations and proofs outlined in the Suppl. Inf.). This composite feature progressively arises in successive rounds of autocatalytic cycles, initially comprising the encounter of the catalyst *E*_1_ provided by a neighboring habitat, with *F*_1_ evolved autocatalytically in the habitat, producing the primordial catalyst *E*_2_ that is capable of amplifying *F*_1_ but not *P*. It is the twofold consequence of the primordial parasite-specificity of *E*_2_, and of the complexity of *P* that exceeds the processivity of the protoreplicase. As a corollary and as displayed in Fig. 2, selection of such molecules confers considerable resistance to parasite invasion, and restores host template and catalyst abundance (Model 1^(0)^ Methods). There is both, strong theoretical and experimental evidence to support these findings. Indeed, while it is generally assumed that catalysts arising in early autocatalytic cycles have low activity and high promiscuity [37], an in depth physico-chemical analysis of how enzymes meet demands of selectivity and accuracy points towards accuracy-rate tradeoffs [38]. Selectivity (accuracy, specificity) implies the ability to discriminate between two templates, provided both are present [37,38], and rate denotes speed [39]. Ground state discrimination means that specificity is achieved mainly through substrate binding, which imposes strong accuracy-rate tradeoffs [38]. Improvements in selectivity mediated by tighter cognate substrate binding invariably lead to lower catalytic efficiency (parallel decreases in the constants for the cognate substrates *K*_*M*_^*cog*^ and *k*_*cat*_^*cog*^; [38]). This is precisely what the model stipulates, the protoreplicase has a primal parasite-selectivity, which comes at the cost of a relatively reduced catalytic activity, yet it allows for adequate host molecule-, but not parasite-amplification. In conclusion, if catalysts with relatively reduced function in the quasispecies confer parasite resistance, yet restore host catalyst and template abundance, they will be positively selected [35].

**Figure 2.**
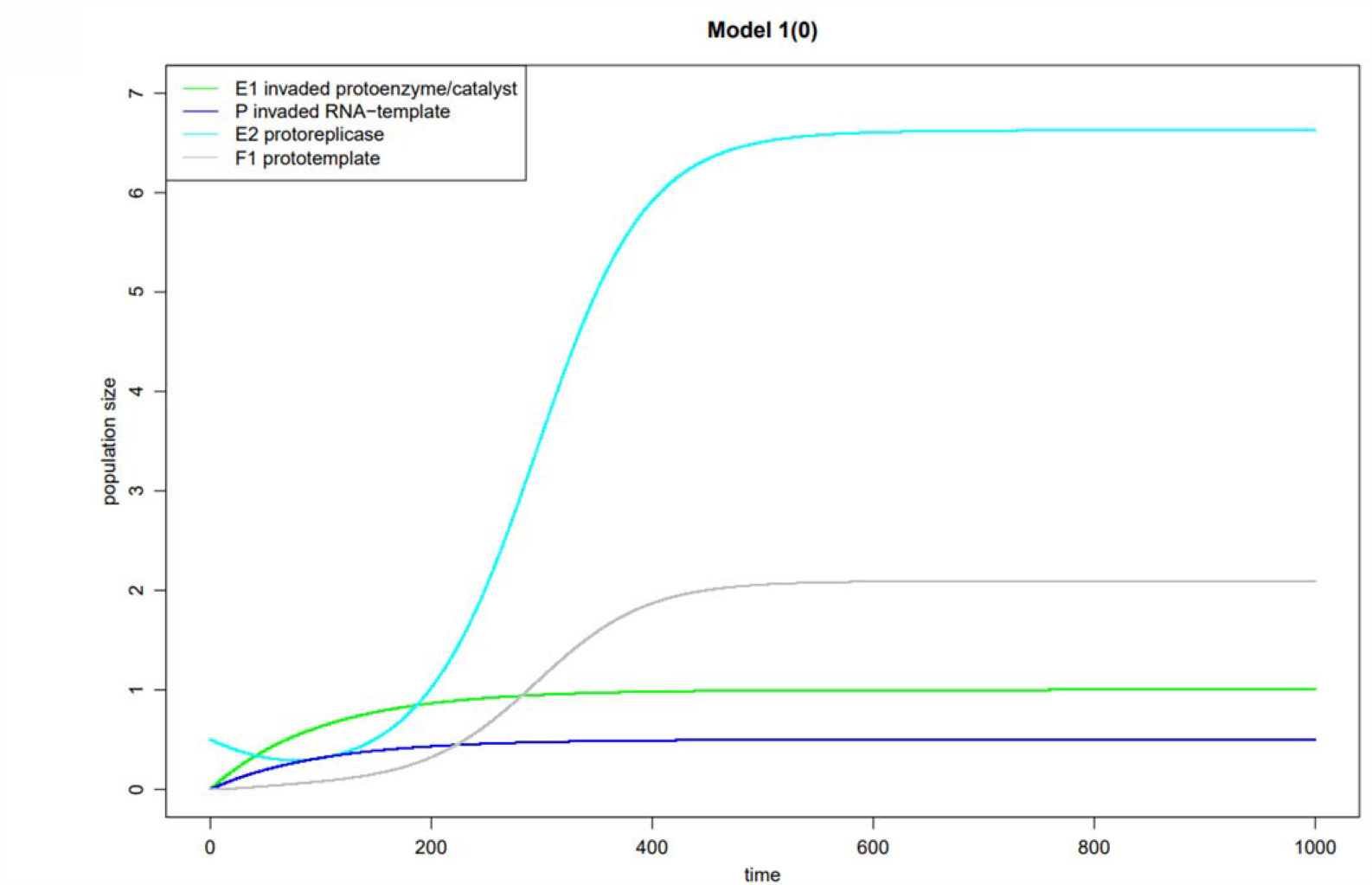
Habitat with primordial autocatalytic cycle. Parasite-specific catalysts with relatively lower catalytic activity in the habitat (*E*_2_ protoreplicase) will confer parasite resistance and thus ensure abundance of habitat molecules in the presence of parasite inflow. Host molecules are expectedly less prevalent compared to Fig. 1B, at least prototemplate *F*_1_.

### Evolution of the primordial autocatalytic cycle into a full-fledged autocatalytic cycle

Natural selection in the mutant quasispecies space will ultimately produce, starting from *E*_2_, a fully efficient, self-replicating ribozyme (replicase *R*_2_) that is bestowed with high catalytic activity [14],capable of generating all enzymes and templates in the habitat (equations and proofs outlined in the Suppl. Inf). However, this has a cost. *R*_2_ is promiscuous, catalytically highly efficient and therefore exposed to parasite invasion, which was not the case for its primordial catalyst precursor with relatively lower activity (protoreplicase *E*_2_, Model 1^(1+)^ stabilized host-parasite-hyperparasite cycles, Methods).

At this point, the protoreplicase *E*_2_-subdomain of *R*_2_ confers *P* selectivity, restoring parasite resistance and rescuing the abundance of the partial catalyst and its templates, even with a moderate parasite inflow (*α*> 0, Fig. 3 left panel). This process has as its central feature a hyperparasite cycle (equations and proofs outlined in the Suppl. Inf.). It is further noteworthy that *F*_*2*_ (formerly *E*_2_) is still produced (Fig. 3), and is acting additively to *R*_2_, which is important to generate antiparasitism later.

**Figure 3.**
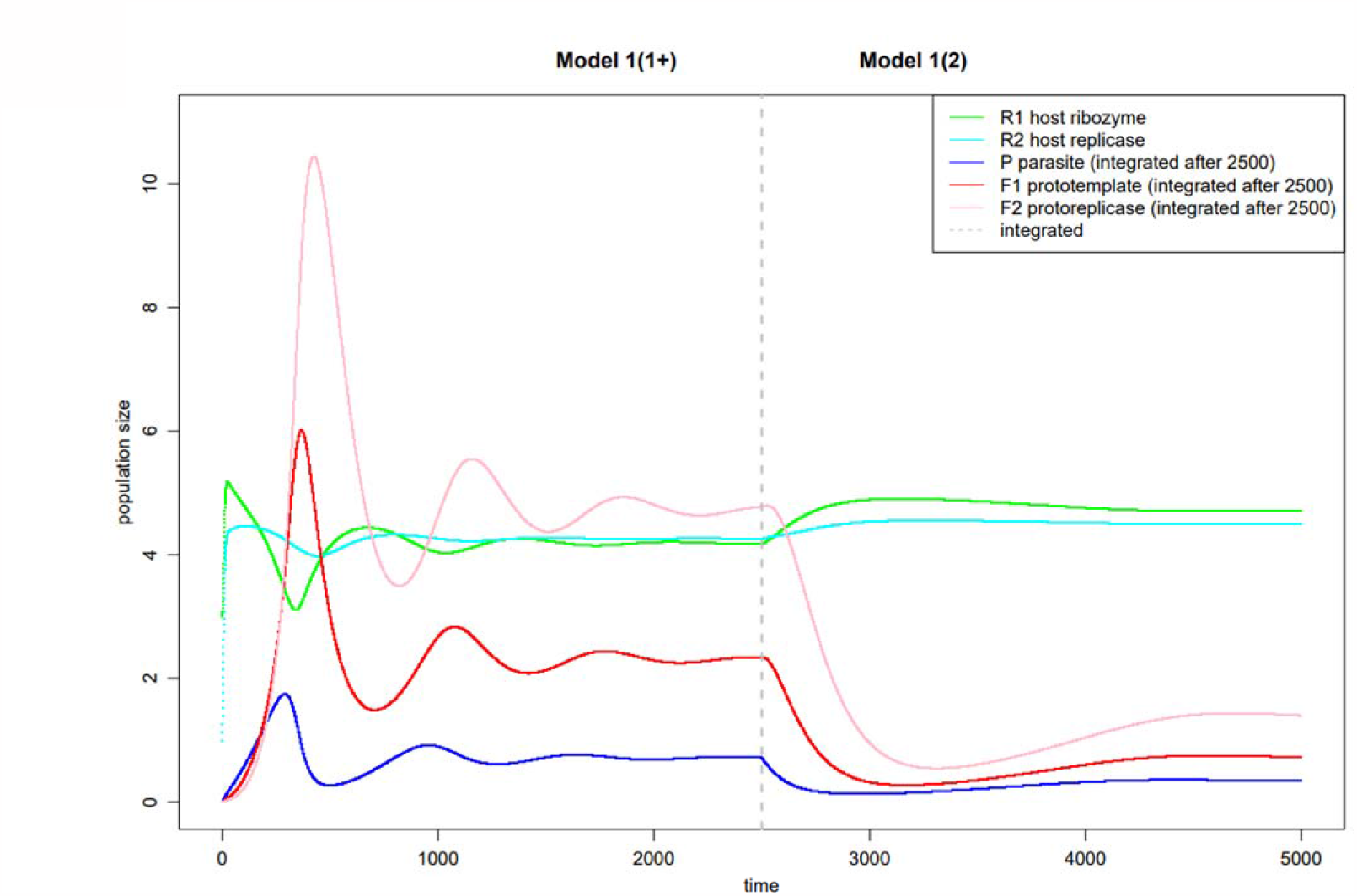
Stabilized host-parasite-hyperparasite cycle with variable parasite inflow. Fig. 3 left panel (Model 1(1+)). A habitat with highly efficient catalysts (self-replicase *R*_2_, template *R*_1_) becomes stabilized with moderate parasite inflow via its relatively less active catalyst-subunit *E*_2_ that confers parasite resistance. *F*_2_ (formerly *E*_2_ in Fig. 2) is generated as well, it acts additively with *R2* and will be important for antiparasitism. Fig 3. right panel (Model 1(2)). Under constant, moderate and thus manageable parasite inflow (*P*^*special*^, Suppl. Inf.), the hyperparasite cycle mutates and becomes integrated into the habitat, even when parasite inflow ceases (prebiotic heredity and mutation-selection balance). Host molecule prevalence is comparable to a habitat without parasites (Fig. 1B).

Under permanent and moderate, thus manageable parasite inflow (*P*^*Special*^, Suppl. Inf.), an increasingly ramified network of autocatalytic cycles arrives in the habitat that persists even when the parasite exposure ceases, constituting a historical record of past parasite exposures (*α* = 0, Fig. 3 right panel).

There is considerable evidence in favor of the essential components of this model, in particular the catalyst accuracy-rate tradeoff and the hyperparasite cycle. Firstly, a solid body of theoretical and experimental evidence supports catalyst accuracy-rate tradeoffs [37,38]. Secondly, metagenomic studies demonstrate the virtually universal presence of tripartite constituents across molecular parasite habitat classes, most persuasively for bacteria-phages-phage satellites [24,25], for RNA viruses partnering the host-virus-defective interfering RNAs [26], and for NCLDVs, with eukaryotic host-NCLDV-virophage [27]. Thirdly, additional support comes from the observation that hyperparasites are induced across the range of parasite concentrations, and especially at high multiplicity infection for RNA viruses [40,41].

### Emergence of antiparasitism in autocatalytic cycles

Under conditions, where the sheer number and complexity of incoming molecular parasite populations would overwhelm the carrying capacity of the habitat, a degenerative process is initiated that erodes catalysis. Formally, the hyperparasite cycle degenerates to an antiparasite cycle (equations and proofs outlined in the Suppl. Inf.; Model 1^(1-)^, Methods). In Figures 4 and 5, the transition from hyperparasitism to antiparasitism is illustrated, in the sense that both coexist at equal activity, partial catalyst and non-catalytic antiparasitic activities (Fig. 4A, see Methods for parameters). In Figure 4B, antiparasitism is fully implemented; hyperparasitism has been fully replaced by the non-catalytic antiparasitism (reproduction parameters hyperparasite *d* and *δ*_3_ are set to zero: *δ*_3_ = *d* = 0, making the difference between pure antiparasitism, and the transition from hyperparasitism to antiparasitism minor. In Figure 4B, the population fluctuations are more volatile than in Figure 4A, but pure antiparasitism does effectively stabilize the autocatalytic cycle, without a hyperparasite component. Figure 5 shows the gradual transition from pure hyperparasitism (top panel) to antiparastism (botton panel), in four successive steps (0, 0.25, 05. 0.75 and 1), progressively replacing hyperparasitism with antiparasitism. Therefore, we conclude that antiparasitism can replace hyperparasitism in situations with a stronger and highly diverse parasite inflow that would overrun the carrying capacity of the autocatalytic cycle in the habitat.

**Figure 4.**
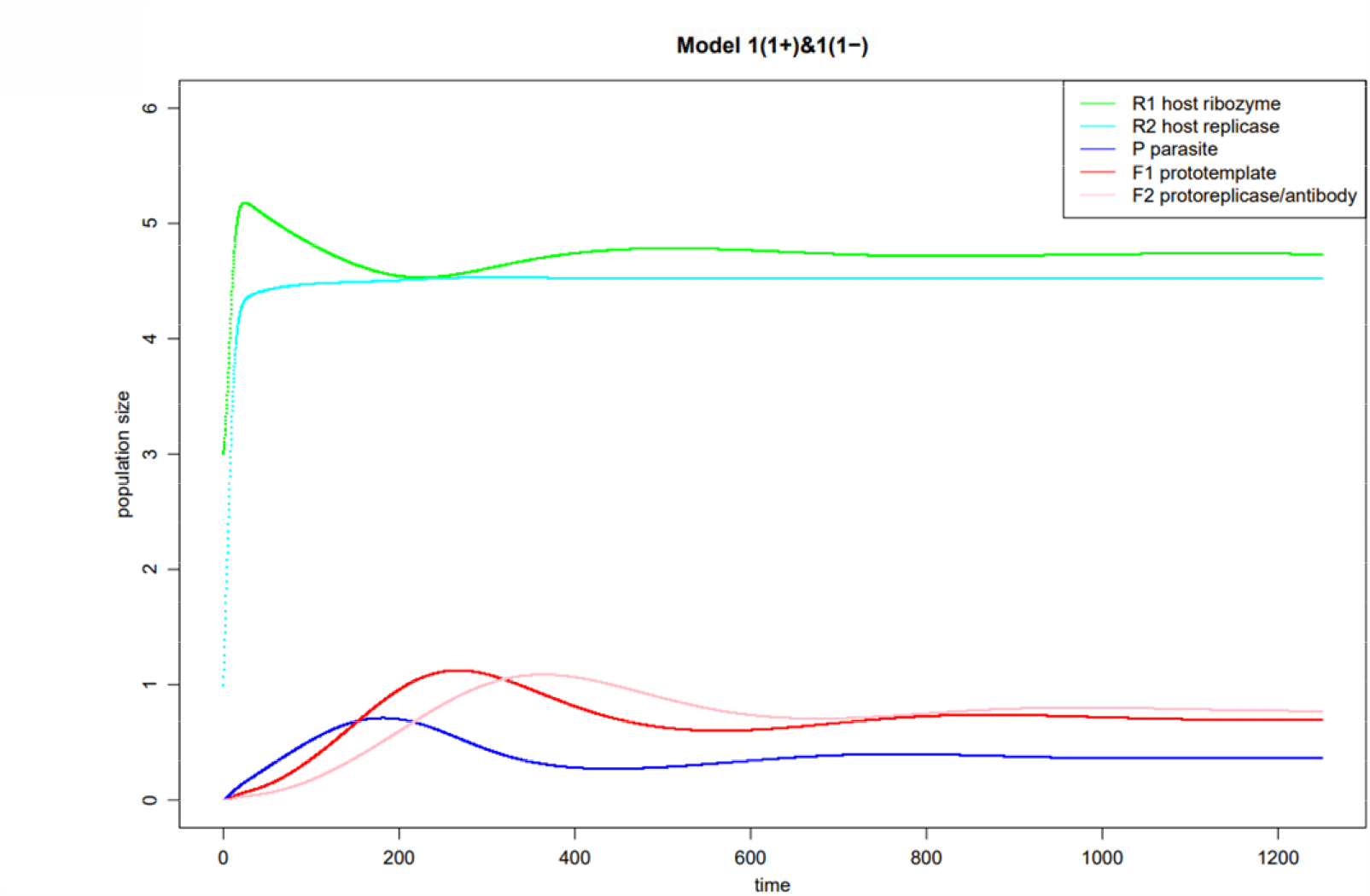
Transition from hyperparasitism to antiparasitism under strong parasite inflow. Fig. 4A Under strong, permanent exposure to complex parasite populations the hyperparasite cyc partially loses its catalytic activity and becomes antiparasitic. The protoreplicase *F*_2_ now continues to function as a partially active catalyst, but also exerts a non-catalytic antiparasitic function (antibody). The parameters in Fig. 4A are set to have these two activities equally represented.

**Fig. 4B.**
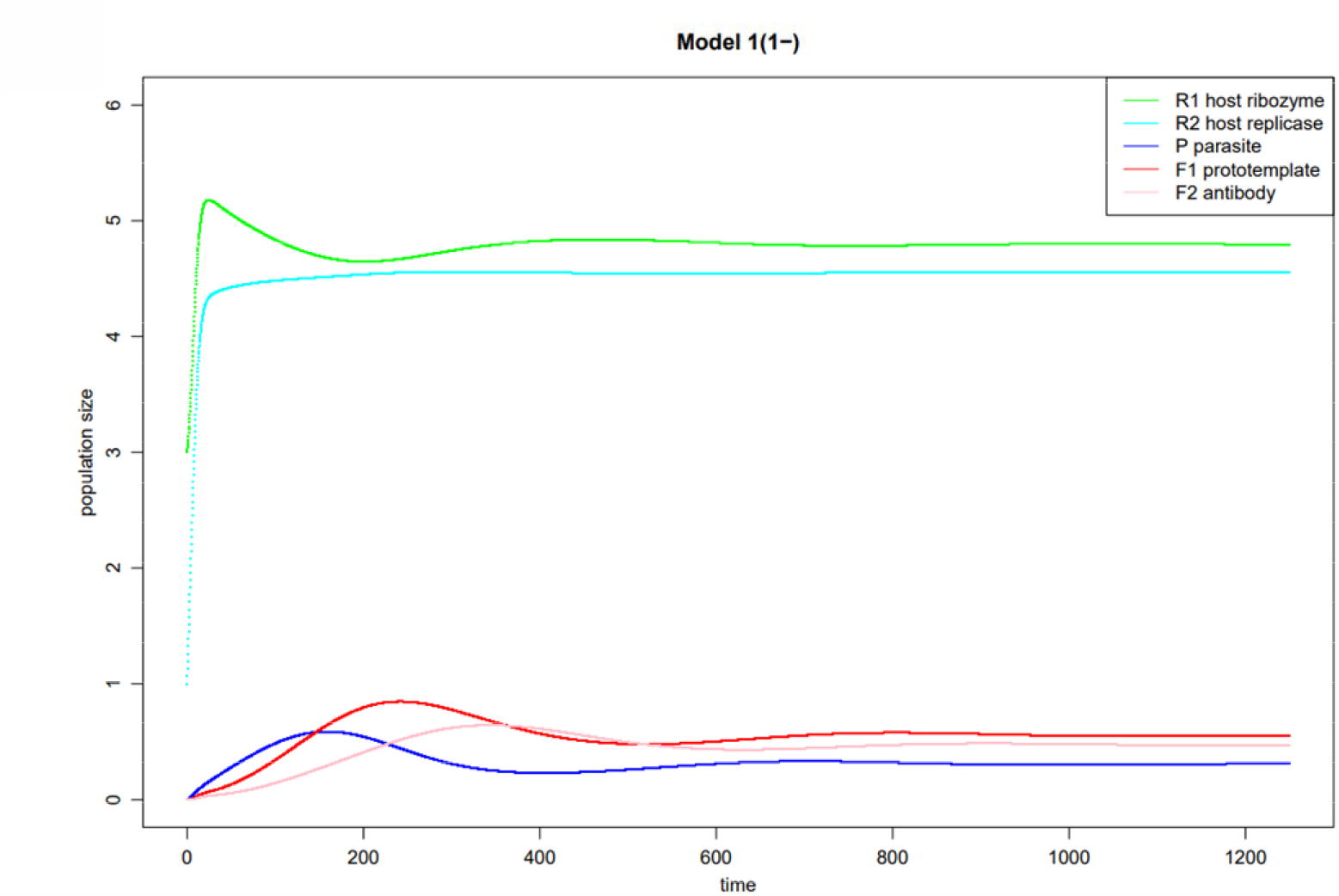
The parameters are now set to completely shift to antiparasitism (fully degenerated hyperparasite cycle, completely non-catalytic). Antiparasitism has fully replaced hyperparasitism, and the habitat is still stabilized, yet with stronger fluctuations.

**Figure 5.**
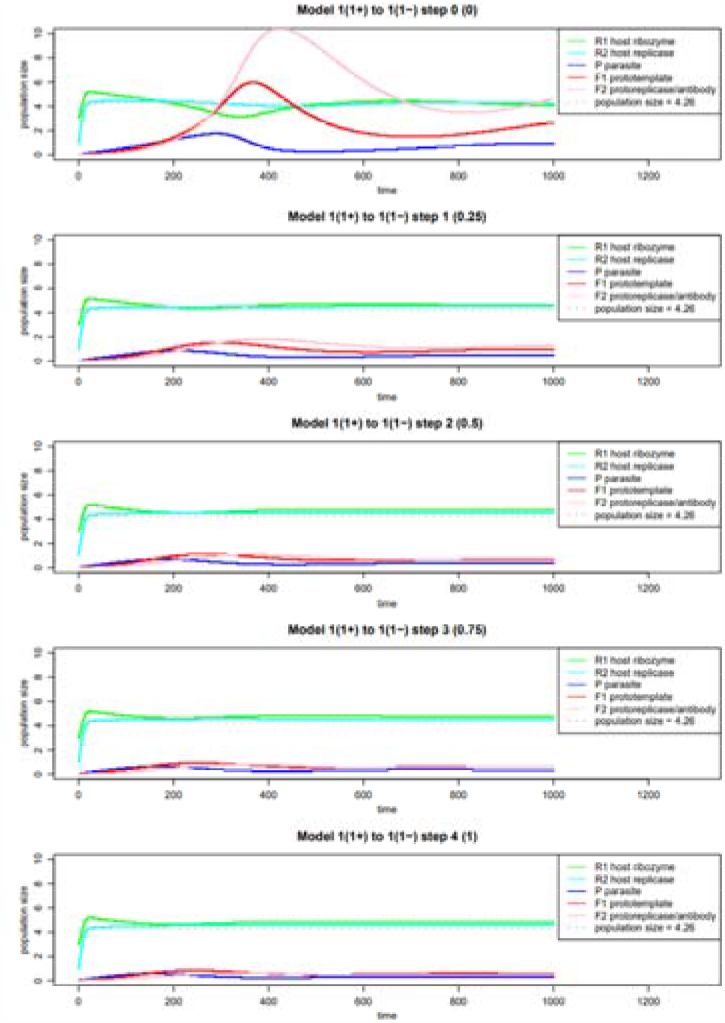
Gradual transition from hyperparasitism to antiparasitism in four steps. The stepwise transition from pure hyperparasitism (top panel) to pure antiparasitism (botton panel) in four steps is shown (0, 0.25. 0.5, 0.75 and 1).

## Discussion

We have outlined in a stepwise fashion, how increasingly elaborate autocatalytic cycles gradually emerge in an early life RNA-habitat. Initially, catalysts with low catalytic activity are selected, because they are parasite-specific and confer resistance (primordial autocatalytic cycles). Soon, complexed mixtures evolve, featuring self-replicating catalysts of high catalytic activity that become stabilized through the selection and integration of hyperparasite cycles, including relatively less active catalyst-subunits that confer parasite resistance (stabilized host-parasite-hyperparasite cycles). Under conditions in which there is the influx of broad and complexed molecular parasite populations into the habitat, the carrying capacity of the autocatalytic networks are exceeded, and a non-catalytic, degenerative process of hyperparasite cycles ensues that leads to the appearance of antiparasitism, and the birth of immunity in early life.

One of the strong features of this model is the solid empirical evidence supporting it. First of all, the catalyst tradeoff analyses [38] applied to autocatalytic RNA networks predicts the emergence of increasingly active ribozymes, with binding promiscuity and high catalytic activity being balanced with specificity and relatively lower activity, which is precisely what the model shows. Prebiotic models indeed support initially fast replicating communities with high molecular diversity, evolving later into more faithful replicators with a narrower repertoire [42]. Moreover, tripartite microbial populations composed of a host, a parasite and a hyperparasite are virtually ubiquitous in the virosphere, as evidenced for bacteria-phages-phage satellites [24,25], host-RNA viruses-defective interfering RNA viruses [26], and eukaryotic host-NCLDVs-virophages [27]. We previously developed a hyperparasite framework based on the LV-equations that corroborates the superior homeostatic viability of tripartite habitats [23]. It thus becomes likely, that early life network habitats with RNA populations were constructed following this architectural logic.

Second, some sort of trans-hierarchical self-organization has to be stipulated at this stage of the early RNA world [43], ideas conceptualized under various designations, such as prebiotic heredity or ecology, and autopoiesis [35,36]. The early RNA world for instance implies lipid-vesicle mediated compartmentalization and supramolecular selection [35]. Applied to our findings, ribozymes conferring parasite resistance are selected early on, survive autopoietically within the quasispecies, and can thus be reselected at later stages, once more efficient catalysts expose the community to waves of threatening parasite invasions.

Third, it is striking that the appearance of microbial immunity is associated with hyperparasites. Indeed, several recent studies show that phages and their satellites encode a set of diverse antiphage systems [24,44], and defective interfering RNAs activate innate immunity [26,40]. Phage satellite-encoded immune systems protect bacteria from phage predation and were proposed to be an integral innate immune component [45]. This notion could easily be extended to omnipresent, bona fide hyperparasites, such as endogenous retroviruses [46,47]. In conclusion, our model provides an overarching theoretical foundation for the notorious association of immunity with the parasite-hyperparasite sphere that is rooted in early life.

Furthermore, it should be noted that extant RNA viral quasispecies most closely resemble the primordial RNA populations described here. Large populations exhibiting fast replication recapitulate the fundamental tradeoff, favoring speed over polymerase accuracy [39]. High mutation rates enable the emergence and maintenance of substantial genetic variation, which leaves marks of the past ecological history within viral genomes via selective sweeps [48]. Moreover, in all contemporary tripartite models the host diverged to cellular life forms carrying polymerases phylogenetically distinct from those prevalent in viruses and selfish genetic elements [49], thus severely limiting genetic exchange. This represented a marked change from the primordial tripartite populations that freely swapped catalysts. The fact that the integration of the virosphere into early RNA world scenarios has been proposed as an important future agenda issue further valorizes our model [43].

Conceptually, our model capitalizes upon one recent and new school of thought in autocatalytic networks, namely the fact that chemical networks behave as ecological species interacting with one another [9]. Since the common underlying physical principle in catalysis and ecological interactions is the mass action law [8], it is not surprising that the biomathematical tools most accurately capturing the quantitative nature of these interactions are also equivalent, namely equations from the LV-family. Probably one of the most significant merits of our model founded in the equivalence of autocatalysis and ecological interactions is the fact that catalytic processes by their very nature can degenerate and become non-catalytic, and that a degenerative process has innovative power. We can summarize the sequence of events as, i) the catalytic process that is inherently error-prone and generates a quasispecies comprising molecular parasites; ii) sequential ecological interactions between host and parasite catalysts stabilize the habitat, forming tripartite ecological communities of host-parasites-hyperparasites; iii) a subgroup of hyperparasite catalysts in such habitats, when overwhelmed with invading parasite waves, can degenerate and create adaptive immunity against those parasites. Thus, in essence, degenerating catalysts create a new, homeostatic function that allows the habitat to adapt.

This model has by design intrinsic limitations, notably the fact that certain conclusions have yet to be experimentally verified [13,14]. Along the same line, it is impossible to infer causation from statistical associations, in particular the pervasive horizontal gene transfer associated with selfish genetic elements blurs the lines. These aspects will be addressed in future studies.

## Methods or Models

### Definitions and Terms

In what follows we assume the following: the Malthusian fitness *r* of a replicating (biochemical) system (or species) is defined by 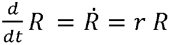, where *R* denotes the population size of the replication sys-tem or species. Evolutionary dynamics between two or more interacting species or systems, e.g. *R,S,P*, … are described by the following type of replicator differential equation e.g. for *R*:

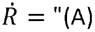 terms of dynamics using habitat resources” x “(B) term of habitat resource restriction” + “(C) terms of dynamics not using habitat resources”

Terms *A* include both “terms of first order kinetics” and “terms of second order kinetics” -“Terms of first order kinetics” are typically natural growth or decay rates, e.g. 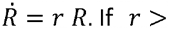. If *r* > 0, growth process described by *rR* eats into resources, whereas if *r* < 0, *r* is a decay rate, and de-cay dynamics does not.

Terms of second order kinetics are derived from encounters between different species. For instance, the term “*bRS*” in 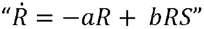 has the following meaning: *bRS* = (*probabiliy of an encounter between members of R and S*) × (*impact of encounter on the fitness of R*). Here we assume, according to the Lotka Volterra calculus that the probability of an encounter between members of *R* and *S* is proportional to the product *RS*.

The term *B* of resource restriction is used in a habitat of limited resources. For instance, in the classic logistic equation 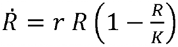 the term 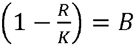 indicates that the habitat has limited resources supporting a maximum population *K* for *R*.

Since every encounter involves resource consuming, we assume here, that all terms of second order kinetics are subject to resource restrictions and therefore need to be multiplied with the term *B*.

Example of a host-parasite interaction (*R*=host, *P*=parasite):

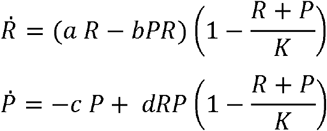

where *a* > 0, *b* > 0, *c* > 0, d > 0. The first order dynamics growth process *aR* and all the second order processes eat into the habitat’s resources, whereas the first order decay process — *c P* does not. The habitat resources restriction is represented by the term 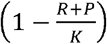 indicating that a maximum population size of *K* = *R* + *P* can be nourished by the habitat and that both, *S* and *P*, compete for the same resources.

### Definition of catalysts, ribozymes and enzymes

We use the term *catalyst*, sometimes also *perfect catalyst*, for any substance that accelerates the rate of a reaction without undergoing any change during the process. We call an *imperfect catalyst* a substance that accelerates the rate of a reaction, but that may undergo a change during the process; antibodies in the transition from hyperparasitism to antiparasitism are examples of imperfect catalysts (see process (3 ′) Suppl. Inf.). *Catalysts* displaying relatively higher substrate specificity generally have relatively lower catalytic activity, for instance the protoreplicase. A subgroup of catalysts is called here enzymes, if they are generated based on RNA templates. An enzyme may be either a *ribozyme*, if it is also used as an RNA template to generate other enzymes, or a *pure enzyme* if it cannot be used as an RNA template for other polymerization processes. The term *ribozyme* encompasses different enzymatic activities, such as recombinase, ligase and polymerase. Recombinases and ligases predominate in early protocycles, while polymerases appear only in advanced networks.

### Model 1^(1)^ habitat with one host autocatalytic cycle exposed to parasite invasion

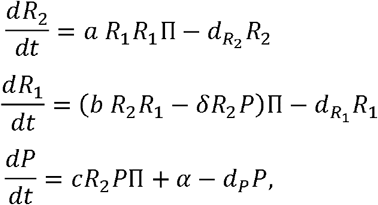

where *a, b* and *c* are positive constants, 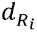, *d*_*P*_ are the decay rates of populations *R*_*i*_, *i* = 1, 2, and of parasite *P* absent its host *R*_2_; *δ* is parasite’s *P* rate of exploitation of the host replicase *R*_2_; and the habitat restriction term is defined as 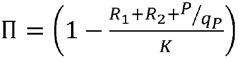 where, *q*_*p*_ > 1, since typically parasites, *P* are more efficient using the resources of the habitat than their hosts *R. α* is the parasite invasion parameter, i.e., per unit of time the habitat is invaded by *α* population units of parasites *P*.

### Model 1^(0)^ habitat with primordial autocatalytic cycle, exposed to template invasion *P* (to become parasite later in evolution) and protoenzyme *E*_1_

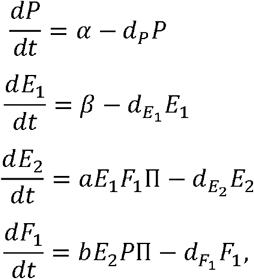

where

*E*_2_ is the protoreplicase produced by this primitive autocatalytic cycle,

*F*_1_ is the prototemplate produced by this primitive autocatalytic cycle,

*α* is the inflow rate of RNA-template *P* into the habitat,

*β* is the inflow rate of protoenzyme/catalyst *E*_1_ into the habitat,

*a* and *b* are positive constants,

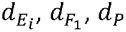 are the decay rates of enzyme populations *E*_*i*_, *i = 1, 2*, of prototemplate 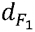, and of RNA-template *P*.

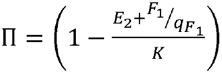 is the habitat restriction term where 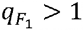, since we assume that proto-templates *F*_1_ are more efficient using the resources of the habitat than the more complex protoreplicase *E*_2_.

### Model 1^(1+)^: stabilized host-parasite-hyperparasite cycle

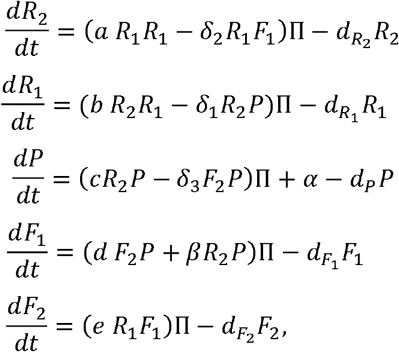

where

*a, b, c, d, e* are positive (reproduction) constants,

*R*_1_ is a ribozyme of the autocatalytic host cycle,

*R*_2_ is the replicase, i.e. another ribozyme (or may be also pure enzyme since it is not used as template) of the autocatalytic host cycle,

*P* is the parasite, exploiting replicase *R*_2_ of the host cycle (exploitation parameter *δ*_1_),

*F*_1_ is the prototemplate exploiting the host *R*_1_ (exploitation parameter *δ*_2_) for protoreplicase *F*_2_ production, triggered upon encounter of *R*_2_ with *P* (trigger parameter *β*) and further produced by protoreplicase *F*_2_ upon encounter with *P*,

*F*_2_ is the protoreplicase (pure enzyme or ribozyme) producing *F*_1_ upon encounter with parasite templates *P* (*F*_*2*_ *= E*_*2*_ from the primitive autocatalytic cycle of Model 1^(0)^), thus, exploiting template of parasite *P* (exploitation parameter *δ*_3_),

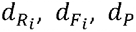 are the decay rates of populations *R*_*i*_, *F*_*i*_, *i* = 1, 2, and of *P*, absent new inflow or reproduction in the habitat,

*α* is the inflow rate of RNA template *P* into the habitat,

*β* is trigger parameter of replicase *R*_2_ producing *F*_1_ by encounter with parasite *P*,

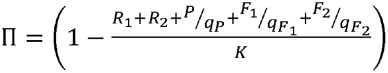 is the habitat restriction term where 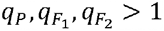, since we assume that parasitizing cycles of *P* and *F*_1_, *F*_2_ are more efficient using the resources of the habitat than the more complex host cycle *R*_1_, *R*_2_.

### Model 1^(2)^:integrated host-parasite-hyperparasite cycle without new parasite inflow

The simulation results of Model 1^(2)^ are realized as follows: Model 1^(1+)^ provides the starting basis, with arbitrary initial populations to establish a stabilized situation, given a moderate parasite inflow with *α* > 0 (Fig. 3 lefthand part). Subsequently, Model 1^(1+)^ is adapted to Model 1^(2)^. All the parameters of Model 1^(2)^ are the same as those of Model 1^(1+),^ with the exception of the parasite inflow, now *α* = 0. As initial conditions for the populations of Model 1^(2)^, we assume the terminal populations of the simulations of Model 1^(2)^, as calculated for Fig. 3 at the beginning of the righthand part. The larger fluctuations of the population at the beginning of the righthand part of Fig. 3 stem from the transition from Model 1^(1+)^ to Model 1^(2)^, where the parasite inflow abruptly decreases from *α* > 0 to *α* = 0. The fact that parasites in Model 1^(2)^ are now reproduced without new inflow shows that they can be viewed as fully integrated into the autocatalytic network, realized in 1^(2)^ (see Suppl. Inf.).

### Model 1^(1−)^: transition from hyperparasitism to antiparasitism

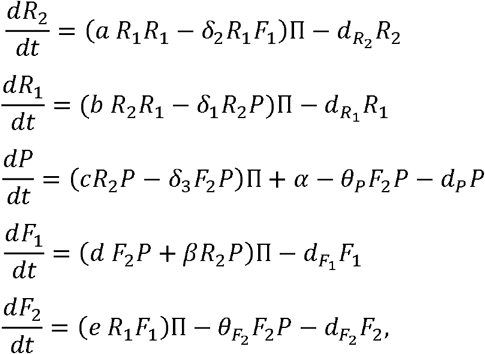

Where all the parameters are the same as in Model 1^(1+)^, with the exception that now two additional parameters are added, namely *θ*_P_ and 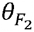. The term *θ*_P_*F*_2_*P* denotes the number of population units of parasites *P* made unviable per unit of time during the encounter of protoreplicase *F*_2_ with *P*, now also functioning as antibody. In addition, the term 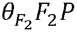 denotes the number of population units of protoreplicase/antibody *F*_2_ made unviable per unit of time during the encounter of protoreplicase/antibody *F*_2_ with *P*. For instance, if 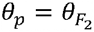, then the same number of antibodies and parasites are made unviable during their encounter. In the evolutionary transition from hyperparasitism to antiparasitism *F*_2_ can still act both ways, as imperfect protoreplicase and as antibody directed against the parasite *P*. We call it *imperfect* protoreplicase or imperfect catalyst, because it still produces/catalyzes prototemplate *F*_2_ upon encounter with template *P*. But now *F*_2_ may be damaged during this process, therefore cannot be called a *perfect* catalyst anymore. The replication differential equations of Model 1^(1−)^ realizing a pure antiparasitism situation are the same as in Model 1^(1−)^ of Fig. 4A, except that the reproduction parameters hyperparasite *d* and δ_3_ are now set to zero: *δ*_3_ *= d* = 0.

## Acknowledgements

We thank Thomas Curran for discussions.

## Supporting information captions

### Self-replicating RNA molecules

Let us consider a set of populations {*R*_1,_ *R*_2,…,_ *R*_*n*_ }of different RNA molecules or enzymes in a primordial RNA setting. Enzymatic functions are exerted by RNA molecules called ribozymes. Ribozymes and other enzymes are forming a catalytic cycle, if

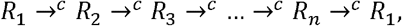

where the symbol “*A*→^c^*B*” means: *A* catalyzes *B*.

In this setting, RNA *A* can display two different functions, the coding and the enzymatic function. RNA-molecules without enzymatic activity have at least a coding potential. Let us call the coding function “template” of RNA molecule. A pure enzyme not bearing a coding function lacks the ability to serve as template. Let us further assume that in this instance successful enzymatic activities require both enzyme and template. An enzymatic activity is thus exerted by a ribozyme or a pure enzyme upon encounter with an RNA template. For example, consider the following equations:

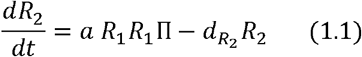

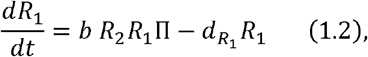

where *a* and *b* are positive constants, 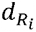 are the decay rates of populations *R*_*i*_, *i*= 1,2 and

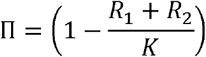

the habitat restriction constant. *K* is the maximum number of *R*_1_ or *R*_2_ molecules, the habitat can feed.

Replication equations (1.1),(1.2) form a two-step autocatalytic cycle. In (1.1),*R*_*1*_ serves both as enzyme and template, in (1.2), *R*_2_ serves as enzyme and *R*_1_ as template, i.e. upon encounter of two different molecules *R*_1_, one serving as template and the other as enzyme, a ribozyme or pure enzyme *R*_2_ is produced/catalyzed, and upon encounter of a molecule *R*_1_ serving as template with a molecule *R*_2_ serving as enzyme the original ribozyme *R*_1_ is produced, forming a complete replication cycle. We symbolize this autocatalytic process by:

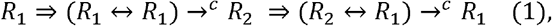

where the symbol “⇒” means “enables” and the symbol “*A*↔*B*” means “encounter of *A* with *B*” and, as already stated above the symbol “→^c^” means “catalyzes”.

Thus, autocatalytic cycles can take the form:

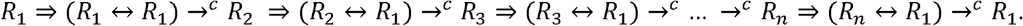

In this example, only *R*_1_ is used as a template during every enzymatic subactivity of the autocatalytic reproduction cycle; every enzymatic activity produces a new ribozyme or a new pure enzyme *R*_*i*_, *i* = 1,… *n*.

Rather than producing a further ribozyme or a pure enzyme, an enzymatic activity can also produce a further template, e.g.

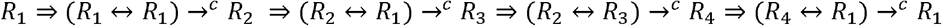

reads as follows: a ribozyme *R*_1_ enables the encounter with another copy of itself, one serving as enzyme, the other as template, and both producing a new enzyme *R*_2_. Upon encounter with a template *R*_1,_ *R*_2_ catalyzes a new template *R*_3_, and upon encounter with this new template a new enzyme *R*_4_, which in turn catalyzes the final reproduction of a new ribozyme molecule *R*_1_ using another molecule *R*_1_ as template.

### Parasitism in autocatalytic cycles

To outline the problem of parasitism, e.g. in the autocatalytic cycle (1), the ribozyme *R*_2_, which may also be a pure enzyme, is acting as a replicase for *R*_1_. Such a replicase is promiscuous and highly active, therefore, *R*_2_ will inevitably be parasitized by parasitic RNA-templates *P*:

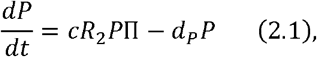

or, written in above introduced technology template format:

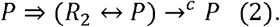

Here again, *c* is a positive constant,*d*_*P*_ > 0 is the decay rate of parasite *P* absent its host *R*_2_, and the habitat restriction term Π extends to

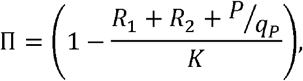

where *q*_*P*_ > 1, since typically parasites *P* are more efficient at using the resources of the habitat compared to their hosts *R*. A ribozyme or pure enzyme molecule *R*_2_ that is used as the replicase for *P* cannot at the same time be used as the replicase for *R*_1_, thus *P* parasitizes the whole autocatalytic cycle (1).

### A stable, primordial autocatalytic cycle (1^origin^)in the presence of parasites

Let us assume the primordial habitat has an abundant inflow of RNA templates *P* of relatively advanced complexity from external sources, and also an abundant inflow of a primary protoenzyme *E*_*1*_. In this mixture, containing *P* and *E*_*1*_, we assume that the following primordial autocatalytic cycle has already evolved. The cycle produces a prototemplate *F*_1_ and a protoreplicase *E*_*2*_ obeying the following autoca-talytic equations:

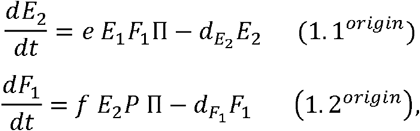

where *e* and *f* are positive constants and 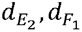are, again decay rates as the ones above. And let the habitat restriction term Π be accordingly defined to fit this situation. Thus, protoenzyme *E*_1_ and temp-lates *P* are given through inflow into the habitat, and ribozyme *E*_2_, which may also be a pure enzyme bearing no template function, and prototemplate *F*_1_ are autocatalytically reproduced.

This original primitive autocatalytic process reads in above format as follows:

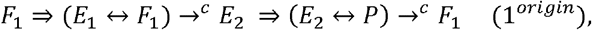

The protoenzyme *E*_1_ produces a protoreplicase *E*_2_ encoded by prototemplate *F*_1_. The fact that *E*_2_ is only a protoreplicase means that is not yet capable of efficiently replicating a complex RNA template P. But provided *P* is above a given threshold complexity, the protoreplicase *E*_2_ can at least produce copies of the prototemplate *F*_1_. As long as the habitat has a minimal threshold concentration of both, acceptably complexed templates *P* and protoenzymes *E*_1_, the archaic autocatalytic cycle (1^*origin*^) can thrive and produce further copies of *F*_1_ and *E*_2_. Remarkably, by design the primitive autocatalytic cycle is immune against parasite invasion by *P* trying to exploit the protoreplicase *E*_2_ for its own replication. On the contrary, the primordial autocatalytic cycle (1^*origin*^) depends on template invasion *P*– in other words parasite invasion – for its own reproduction.

### Evolution of the primordial autocatalytic cycle (1^*origin*^) into a full-fledged autocatalytic cycle(1)

We surmise that the host cycle (1) is capable of prebiotic heredity [10-12], that the primordial autocatalytic cycle (1^*origin*^) has been conserved in the quasispecies, and therefore can again be selected for upon parasite invasion. Therefore, the protoreplicase *E*_2_ can be reselected from the quasispecies. This triggered process can take the following form:

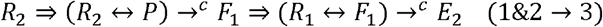

The fully functional replicase *R*_2_ is designed to replicate *R*_1_ according to (1.2),but it can also be exploited to replicate genetic parasites *P* according to (2). Upon encounter with a parasitic template *P*, it not only replicates *P* but also produces via its catalyst subunit *E*_2_ a prototemplate *F*_1_, as did its parent *E*_2_ replicase. What is true for the replicase *R*_2_ – that it is more efficient than its parent catalyst, the protoreplicase *R*_2_ – is also true for ribozyme *R*_1_. It is catalytically more efficient than protoenzyme *E*_1_ it had replaced during evolution. *R*_1_ can not only produce a full-fledged replicase *R*_2_ given another copy of itself as a template, but via its protoenzyme *E*_1_-subunit it reproduces the protoreplicase *E*_2_ upon encounter with prototemplate*F*_1;_

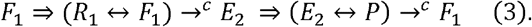

In order to name the processes:

1. is the host cycle
2. is the parasite cycle exploiting the host cycle
3. is the hyperparasite cycle, parasitizing the host (exploiting ribozyme *R*_1_ for protoreplicase *E*_2_ production), and parasitizing the parasite *P* (exploiting its template for prototemplate *F*_1_ production) triggered throughout the process(1&2→3).

### Prebiotic heredity of autocatalytic cycles to historical parasite exposure

In an environment with a constant – i.e. moderate, thus manageable – inflow of invasive parasites the full-fledged autocatalytic host-parasite-hyperparasite cycle (1),(2),(3) together with the trigger process (1&2→3) becomes increasingly integrated. If the nature of the constant inflow is always of the same special type of invasive parasites *P*^*special*^, then the hyperparasite process (3) as well as the trigger process (1&2→3)can mutate/adapt to processes that are fully adapted to *P*^*special*^ :

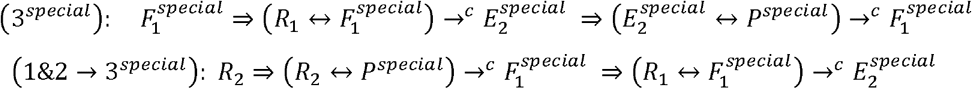

It is reasonable to assume that the host-parasite-hyperparasite process (1),(2),(3) together with the trigger process (1&2→3)mutates under permanent special parasite inflow *P*^*special*^ to an adapted process (1),(2),(3 ^*special*^), together with the adapted trigger process(1&2→3^*special*^) . This network of newly formed autocatalytic cycles reproduces all its elements. All adapted elements *R*_1_, *R*_2_,*P* ^*special*^, 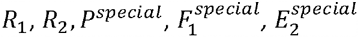 reproduce, even when the special external parasites *P* ^special^ cease to invade the habitat.

With the above we now can say: the primordial autocatalytic cycle (1)^(0)^ has mutated to the cycle (1)^(1)^, which in turn mutates to the cycle (1)^(2)^, after having faced a longer term invasion of the same type of parasites *P*^*special*^ .

With this setup evolution exhibits two salient properties:

- The systems (1)^(0)^, (1)^(1)^, and (1)^(2)^ can be viewed as increasing complexed networks of autonomous autocatalytic cycles. They can be seen as the first steps in kickstarting evolution towards complexed living systems.
- Prebiotic heredity [10-12] allows the cycle-network to preserve the complete record of its historical environmental exposure to parasites:
  ° (1)^(2)^ preserves the record of (1)^(1)^, especially the type of its infection *P*^*special*^, and
  ° (1)^(1)^ preserves the record of its origin (1)^(0)^ being immune against a broad range of para-site infections *P*, and
  ° (1)^(0)^ preserves the record of immunity against a broad range of parasite infections *P*.

As evolution progresses, the autocatalytic cycles (1)^(2)^ and (1)^(1)^ can now be invaded by new parasites, which can be integrated into more complexed systems, e.g. (1)^(3)^ or (1)^(2′)^, which in turn always keep track of the entire historical parasite exposure.

### Emergence of antiparasitism in autocatalytic cycles

Let us assume the autocatalytic cycle (1)^(1)^ is permanently invaded by many different types of parasites *P*, to which it was not yet been exposed historically. The integration of every special subtype of these parasites would not be viable and lead to a complexity overflow in the evolving autocatalytic network. As first line of defense, the emergence of the autocatalytic host-parasite-hyperparasite cycle (1)^(1+)^ = { (1),(2),(3), (1&2→3)}, as previously described would be able to stabilize the situation, if infection *P* is not too overwhelming, and as time advances not too diverse. For a more sustained, overwhelming and diverse parasite attack, another type of immunity would be adapted, although it also originates from the already established host-parasite-hyperparasite stabilization (1)^(1+)^:

In the original primitive autocatalytic cycle (1^*origin*^), *F*_*1*_ ⇒ (*E*_*1*_ ↔ *F*_*1*_)→^*c*^ *E*_*2*_ ⇒(*E*_*2*_↔ *P)* →^*c*^ *F*_*1*_,it is essential the both encounters (*E*_1_↔ *F*_1_) and *E*_2_ ↔ *P* preserve their catalytic nature during the whole process, since otherwise – e.g. if they are damaged, this will heavily reduce their respective population sizes – the autocatalytic nature of (1^*origin*^) could break down rapidly. However, this is not the case for a full-fledged autocatalytic host-parasite-hyperparasite trio (1),(2),(3) in ^(1+)^ that has already evolved. The hyperparasite (3), *F*_*1*_ ⇒ (*R*_*1*_ ↔ *F*_*1*_)→^*c*^ *E*_*2*_ ⇒(*E*_*2*_↔ *P)* →^*c*^ *F*_*1*_ represents the host’s (1) evolutionary heredity from its primordial origin (1^*origin*^). If in this hyperparasite cycle (3) the encounter of e.g. the protoenzyme *E*_2_ with the genetic parasite *P* loses its catalytic nature, then the autocatalytic nature of the full-fledged cycle (1),(2),(3) in (1)^(1+)^ would not break down. If *P*, or both *P* and *E*_2_ are damaged during their encounter, this would only lead to a strengthening of the host cycle (1) which is viable also in the absence of parasite and hyperparasite cycle (2),(3). As a result, the cycles (2),(3) would not break down, given the fact that the habitat still faces a sustained genetic parasite invasion *P*. Thus, the full-fledged cycle (1),(2),(3)would easily survive if the encounter of the protoreplicase *E*^2^ with the genetic parasite *P* would lose its catalytic nature. On the contrary, if *P* would be damaged, the stability, i.e. the fitness of (1)^(1)^ and with it (1)^(1+)^,would increase further. Under such circumstances, we expect evolution to establish a degenerative process of the (*E*_*2*_ ↔ *P*) encounter. We write for such a degenerated encounter:

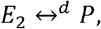

meaning that during the encounter of *E*_2_ with *P*, both elements, *E*_2_ and *P*, or at least the template *P*, are damaged and made unviable:

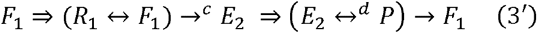

Here the symbol “→” means “produces”. But this “production” is not catalytic anymore, since the “producers” are damaged during the process. For an illustration of this situation where hyper-parasitism coexists with antiparasitism (i.e. *F*_1_ production upon encounter of *E*_2_ with *P* exploiting *P* as template) see Fig. 4A and Fig. 5.

The antiparasite process can now degenerate even further to(3′ ′), where the encounter of prototemplate *E*_2_, which we now call antibodies, with parasite *P* loses its ability to produce the prototemplate*F*_1_:

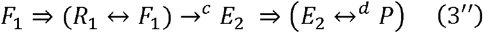

The ability of the antibody *E*_2_ to produce a prototemplate *F*_1_ upon encounter with the parasite template *P* is not an evolutionary necessary one in this situation. Prototemplates *F*_1_ are only needed as long as parasites *P* exploit replicase *R*_2_ for reproduction, but replicase *R*_2_ already triggers *F*_1_-production upon encounter with parasites *P* through the trigger process (1&2→3). Therefore, *F*_1_-production through the antiparasite process (3′ ′)becomes redundant.

## Author contributions

Conception and design of the work: MP and BC. Data acquisition MP; data analysis MP, CI, BC; interpretation of data MP, CI, JAC, BC. Creation of new software used in the work: MP, CI. Drafted the work and revised it: BC, MP and JAC.

## Data availability

The data, including scripts, is fully made available at the following sites (https://github.com/BICC-UNIL-EPFL/parasite-immunity-from-hyperparasitism [github.com]; https://doi.org/10.5281/zenodo.7970788 [doi.org]).

